# Efficacy of an oral lipid nanocrystal (LNC) formulation of amphotericin B (MAT2203) in the neutropenic mouse model of pulmonary mucormycosis

**DOI:** 10.1101/2023.11.22.568278

**Authors:** Yiyou Gu, Teclegiorgis Gebremariam, Sondus Alkhazraji, Eman Youssef, Sabrina El-Gamal, Theresa Matkovits, Jenel Cobb, Raphael Mannino, Ashraf S. Ibrahim

## Abstract

Invasive mucormycosis (IM) is associated with high mortality and morbidity and commonly afflicts patients with weakened immune systems. MAT2203 is an orally administered lipid nanocrystal (LNC) formulation of amphotericin B, which has been shown to be safe and effective against other fungal infections. We sought to compare the efficacy of MAT2203 to liposomal amphotericin B (LAMB) treatment in a neutropenic mouse model of IM due to *R. arrhizus* var. *delemar* or *Mucor circinelloides f. jenssenii* DI15-131. Treatment with placebo (diluent control), oral MAT2203 administered as BID and QD or intravenous LAMB for 4 days, began 16 h post infection and continued for 7 and 4 days, respectively. Survival through Day +21 and tissue fungal burden of lung or brain in animals euthanized on Day +4 served as a primary and secondary endpoint, respectively. In both infection types, MAT2203 was as effective as LAMB in prolonging median survival time (MST) and enhancing overall survival *vs.* placebo-treated mice (*P*<0.05 by Log-Rank). Furthermore, both MAT2203 and LAMB treatment resulted in significant ∼1.0-1.5-log reduction and ∼2.0-2.2-log *in R. delemar* or *M. circinelloides* lung and brain burden, *vs.* placebo mice, respectively. These results support the potential efficacy of oral MAT2203 as an alternative to LAMB. Continued investigation and development of this novel oral formulation of the amphotericin B for the treatment of mucormycosis is warranted.

## Introduction

Diabetic ketoacidosis (DKA), neutropenia, and high dose corticosteroid treatment are all clinical factors that predispose patients for mucormycosis, a life-threatening fungal infection with high mortality rates of >50% (1–3). In the most severe circumstances, patients with brain involvement, persistent neutropenia, and hematogenously disseminated disease have mortality rates >90% (4, 5). Furthermore, COVID-19-associated mucormycosis (CAM) has been reported in many countries during the second wave of the COVID-19 pandemic with > 50,000 cases reported in India between May-August of 2021 (6, 7). Mucorales fungi are the responsible organism, with *Rhizopus* species being the most common cause of infection worldwide followed by *Mucor* species (8, 9).

There are three drugs to treat mucormycosis: amphotericin B-based compounds, isavuconazole, and posaconazole. Lipid formulation of amphotericin B (including liposomal amphotericin B [LAMB]) is the first-line therapy for mucormycosis, while the two azoles are reserved for stepdown and/or salvage therapy (10). Although LAMB is less toxic than amphotericin B deoxycholate, considerable toxicity and inability to deliver the drug orally limits its use and contribute to the high mortality seen with mucormycosis. A recently developed oral formulation of amphotericin B (MAT2203, Matinas BioPharma) using a proprietary lipid nanocrystal (LNC) delivery platform has been shown to be safe when given at doses as high as 40 mg/kg and has demonstrated efficacy against murine aspergillosis and murine cryptococcal meningoencephalitis (11, 12), as well as in a human Phase 2 study or cryptococcal meningitis (13). We sought to assess the activity of MAT2203 in our established immunosuppressed murine models of mucormycosis infected by *R. arrhizus* var. *delemar,* or *M. circinelloides f. jenssenii.* We compared overall survival, median survival time (MST), tissue fungal burden and histology of target organs to those treated with LAMB.

## Results

### Susceptibility testing

We determined the minimum inhibitory concentrations that resulted in 50% (MIC_50_) or 90% (MIC_90_) inhibition of the growth of the fungal spores when compared to no treatment (Table 1). MAT2203 showed 5-10-fold increase in *in vitro* activity against the *R. arrhizus* var. *delemar* and *M. circineloides* when compared to LAMB.

**Table 1:** MIC_50_ and MIC_90_ are defined as the drug concentration that causes 50% and 90% reduction in growth, respectively.

| Drug | *R. arrhizus* var. *delemar* 99-880 |  | *M. circinelloides* f. *jenssenii* DI15-131 |  |
| --- | --- | --- | --- | --- |
|  | MIC<sub>50</sub><br>(µg/mL) | MIC<sub>90</sub><br>(µg/mL) | MIC<sub>50</sub><br>(µg/mL) | MIC<sub>90</sub><br>(µg/mL) |
| LAMB | 0.0156 | 0.0315 | 0.0315 | 0.0625 |
| MAT2203 | 0.0016 | 0.0081 | 0.0039 | 0.0078 |

### The *in vivo* activity of MAT2203 against *R. arrhizus* var. *delemar*

#### Survival

Two survival experiments were conducted to determine the activity of MAT2203 in treating murine mucormycosis due to *R. arrhizus* var. *delemar* 99-880. In the first experiment, neutropenic mice were infected and treated as detailed above. MAT2203 doses of 5 and 15 mg/kg qd showed enhanced overall survival of 20% and 40% when compared to 0% of placebo mice (infection no treatment), respectively. A high dose of 45 mg/kg of MAT2203 did not show benefit over placebo treated mice. Concordant with previous studies (14, 15), LAMB treatment resulted in a 40% overall survival of mice (Fig 1A). A repeat study was conducted to confirm the results and to investigate if twice daily treatment with MAT2203 will result in an enhanced benefit in survival of mice infected with *R. arrhizus* var. *delemar.* MAT2203 at 15 mg/kg given (qd) and LAMB (10 mg/kg, qd) resulted in a similar protection and an overall survival of immunosuppressed mice of 40% and 50%, respectively (Fig 1B). In addition, splitting the daily dose of 15 mg/kg into two doses of 7.5 mg/kg/day (bid) resulted in a similar overall survival of 30% when compared to 15 mg/kg qd dosing. However, treating mice with two doses of 15 mg/kg/day did not protect mice from infection when compared to placebo-treated mice which had 0% survival by day 18 post infection. Collectively, these results confirm the equal protection afforded by oral MAT2203 when given in doses of 5-15 mg/kg qd to the standard of care of LAMB treatment and that daily doses of MAT2203 above 15 mg/kg is not protective.

**Figure 1:**
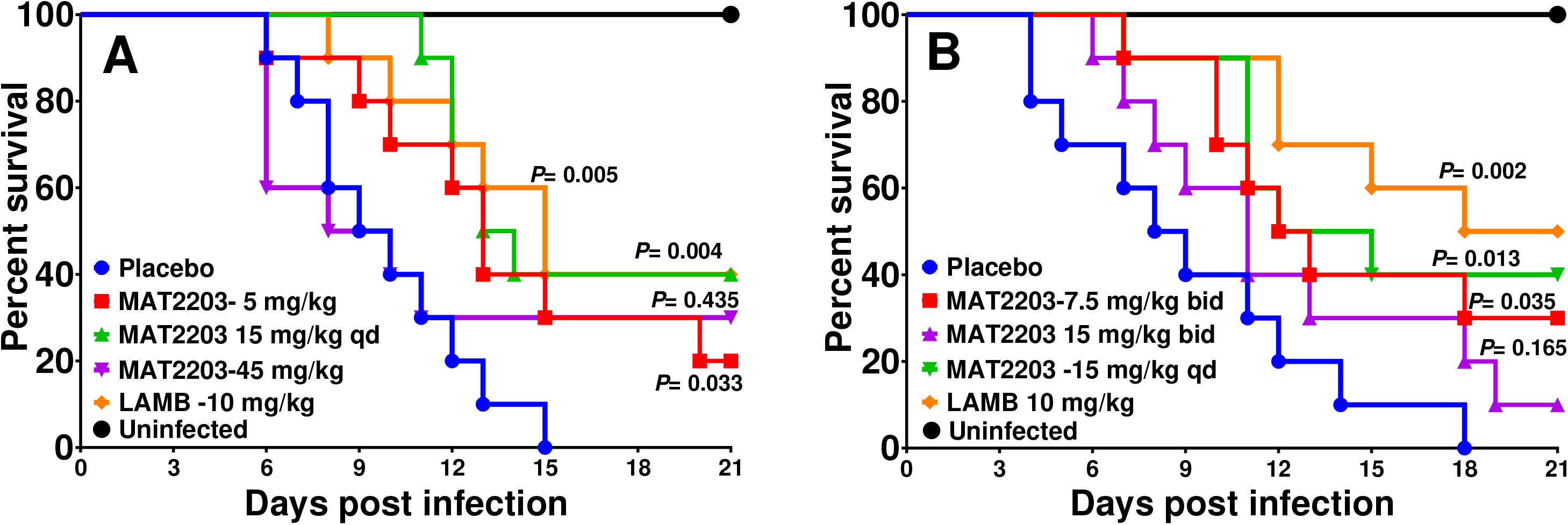
Survival of immunosuppressed mice (n=10/group) infected with *R. arrhizus* var. *delemar* and treated with either MAT2203 or LAMB. Two independent experiments were conducted (Exp 1 [A], and Exp 2 [B]). In experiment 1, The delivered infectious inoculum was 1.8 × 10^4^, and 1.5 × 10^4^ spores for experiment 1 and 2, respectively. Treatment started 16 h post infection and continued for 7 days for MAT2203 and 4 days for LAMB. *P* values on each of graphs are versus placebo-treated mice.

Because of the concordant results in protection seen with the qd dosing of 15 mg/kg of MAT2203 and 10 mg/kg of LAMB, we combined the data of experiment 1 and 2 and measured the MST for each treatment. The MST was concordant with the overall survival by day 21 post infection with doses of 5 and 15 mg/kg qd or 7.5 mg/kg of MAT2203 and 10 mg/kg of LAMB showing improved MST of 13, 13.5, 12.5, and 16.5 days, respectively *vs.* 9 days for placebo-treated mice (Fig 2).

**Figure 2:**
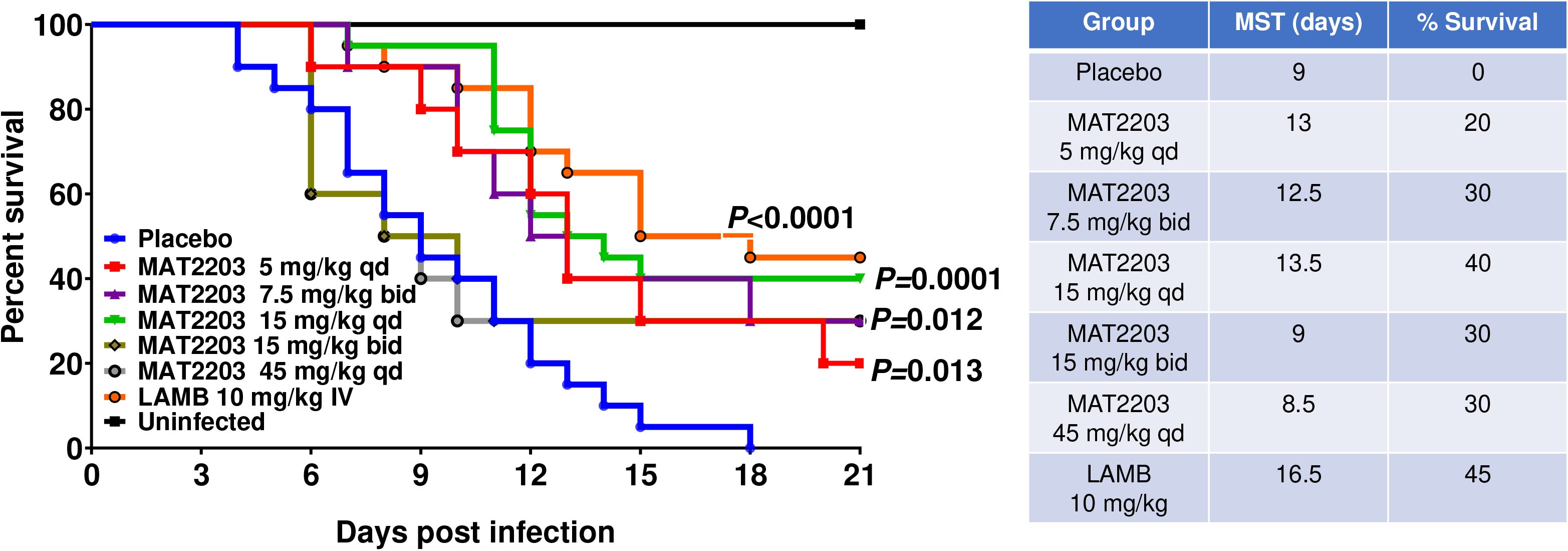
Combined data of survival of neutropenic mice infected with *R. arrhizus* var. *delemar* and treated with MAT2203 or LAMB. *P* values on each of graphs are versus placebo-treated mice. Data in the Table include the median survival times and the overall survival by day 21 post infection.

#### Tissue fungal burden

Because MAT2203 enhanced survival of mice infected with *R. arrhizus* var. *delemar,* we evaluated the effects this drug on the tissue fungal burden of target organs of lung and brain (14, 16, 17). Treating mice with 15 mg/kg of MAT2203 qd resulted in ∼1.5-log reduction in lung and 1.0-log reduction in brain fungal spores when compared to placebo-treated mice. Importantly, this reduction in fungal burden was comparable to reduction seen in mice treated with LAMB. While a dose of MAT2203 at 5 mg/kg once daily trended to lower fungal burden in lung, this difference was not significant (Fig 3).

**Figure 3:**
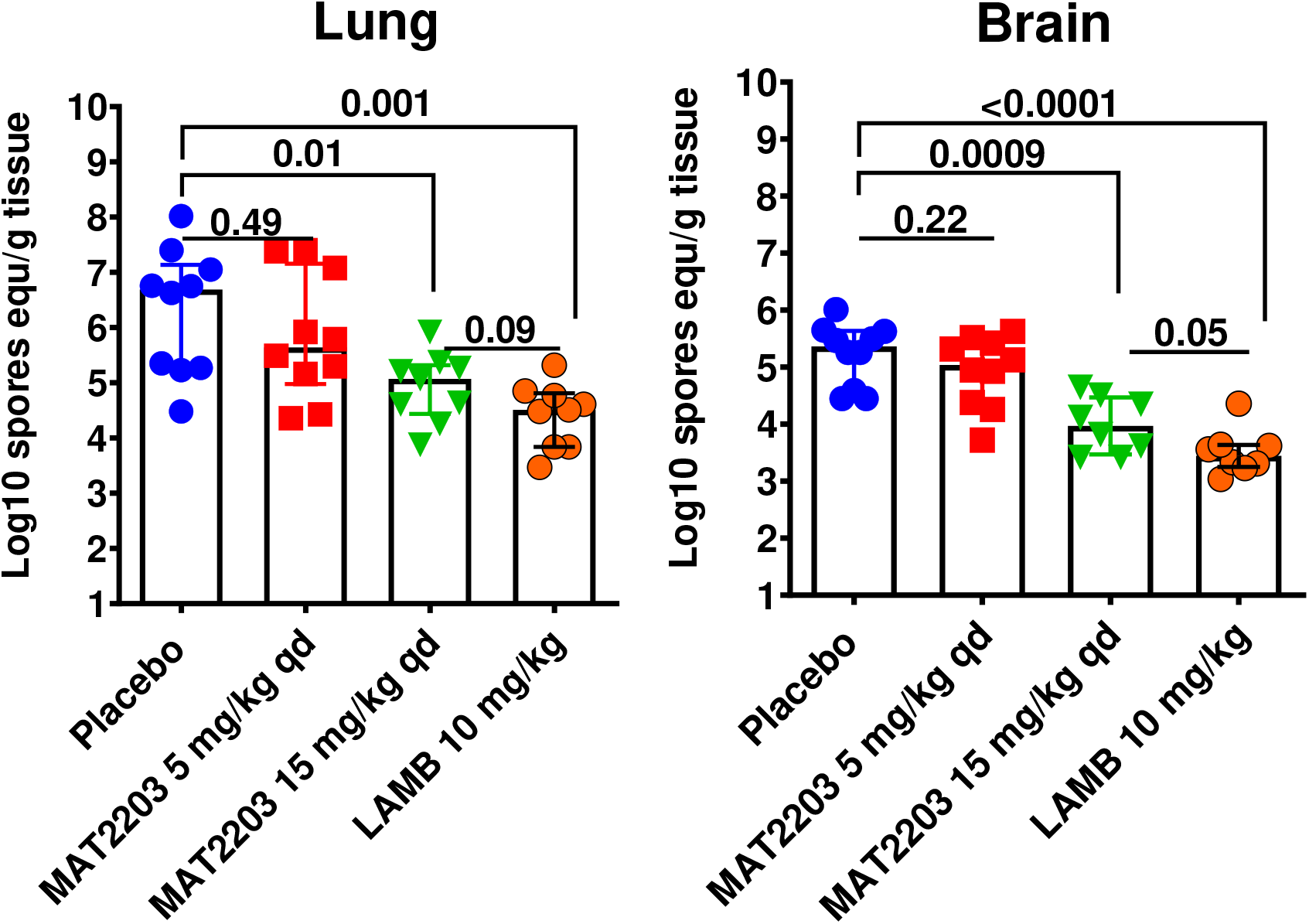
Reduction in tissue fungal burden of immunosuppressed mice infected with *R. arrhizu* var. *delemar*. Mice (n=10/group) infected intratracheally with *R. arrhizus* var. *delemar* (inhaled inoculum of 2.9 × 10^4^ spores/mouse) and 16 h later treated with MAT2203 5mg/kg qd, or 15 mg/kg qd, or with LAMB 10 mg/kg. On day +4 organs were collected and processed for tissue fungal burden by qPCR. Data= median ± interquartile range and the *y* axis represents the lower limit of detection. Intergroup *P* values shown as a dark line. Both MAT2203 at 15 mg/kg and LAMB resulted in a statistically significant reduction in lung and brain fungal burden *vs.* placebo control. (Wilcoxon Rank sum test).

### The in vivo activity of MAT2203 against M. circinelloides f. jenssenii

#### Survival

To investigate if the protective effect of MAT2203 can be expanded to other Mucorales fungi that are commonly isolated from patients with mucormycosis, we infected immunosuppressed mice with *M. circinelloides f. jenssenii* and 16 h later treated them with either MAT2203 (15 mg/kg qd), or LAMB (10 mg/kg qd) as detailed above. Consistent with the results obtained with mice infected with *R. arrhizus* var. *delemar,* both drugs prolonged median survival time by 13.5 and >21 days for MAT2203 and LAMB, respectively, when compared to 5 days of placebo control. Moreover, both drugs enhanced overall 21-day survival to 50% and 60% of MAT2203- and LAMB-treated mice *vs.* 0% survival for placebo control mice (Fig 4).

**Figure. 4.**
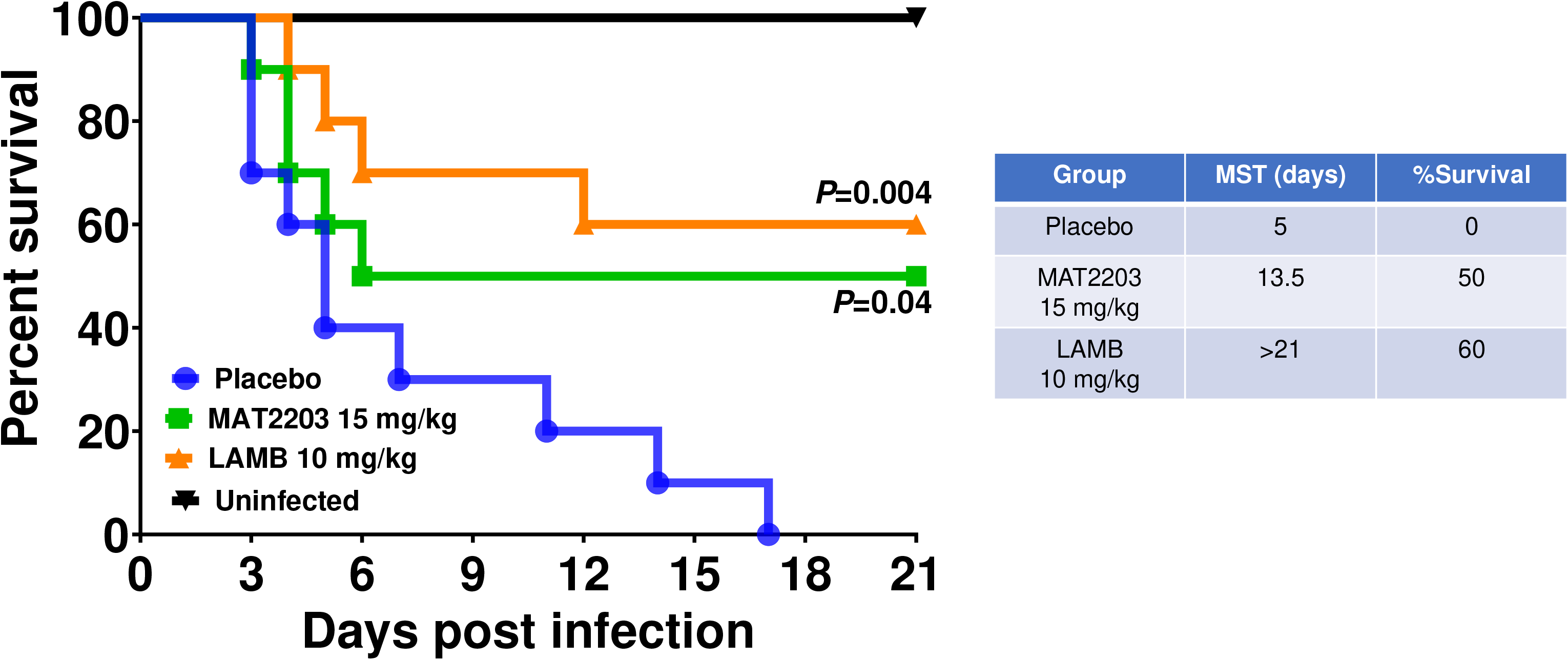
Survival of neutropenic mice (n=10/goup) infected with *M. circinelloides f. jenssenii* and treated with MAT2203 or LAMB. *P* values on each of graphs are versus placebo-treated mice. Data in the Table include the median survival times and the overall survival by day 21 post infection.

**Figure 5.**
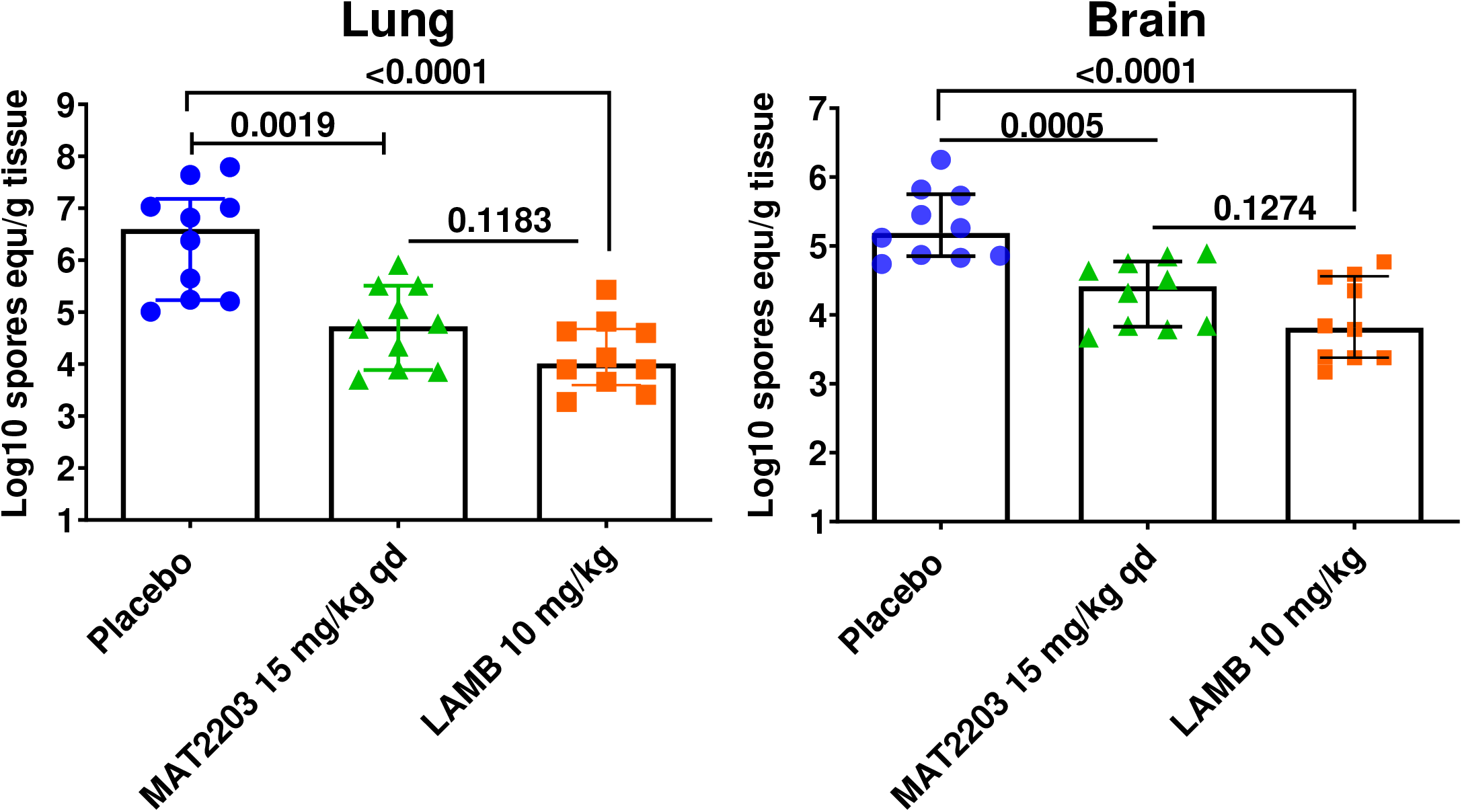
Reduction in tissue fungal burden of neutropenic mice infected with *M. circinelloides f. jenssenii*. Mice (n=10/group) infected intratracheally with 2.9 × 10^4^ spores/mouse) and 16 h later treated with MAT2203 (15 mg/kg qd), or with LAMB (10 mg/kg). On day +4 organs were collected and processed for tissue fungal burden by qPCR. Data= median ± interquartile range and the *y* axis represents the lower limit of detection. Intergroup *P* values shown as a dark line. Both MAT2203 at 15 mg/kg and LAMB resulted in a statistically significant reduction in lung and brain fungal burden *vs.* placebo control. (Wilcoxon Rank sum test).

#### Tissue fungal burden and histopathological examination

Because MAT2203 enhanced survival of mice infected with *M. circinelloides f. jenssenii,* we evaluated the effect of this drug on the tissue fungal burden of target organs of lung and brain (14, 16, 17). Treating mice with 15 mg/kg of MAT2203 qd resulted in ∼2.0-log reduction in the lung and ∼1.0-log reduction in the brain fungal burden when compared to placebo-treated mice. Importantly, this reduction in fungal spores was comparable to reduction seen in mice treated with LAMB.

We also conducted histopathological examination on the same organs processed for the tissue fungal burden experiment. While placebo mice had abscesses full of intact broad aseptate fungal hyphae (consistent with mucormycosis (18)), mice treated with MAT2203 or LAMB had less fungal abscesses with damaged and shorter hyphae (Fig. 6). Collectively, these results demonstrate the equivalent efficacy of MAT2203 to the current standard of care of LAMB.

**Figure 6.**
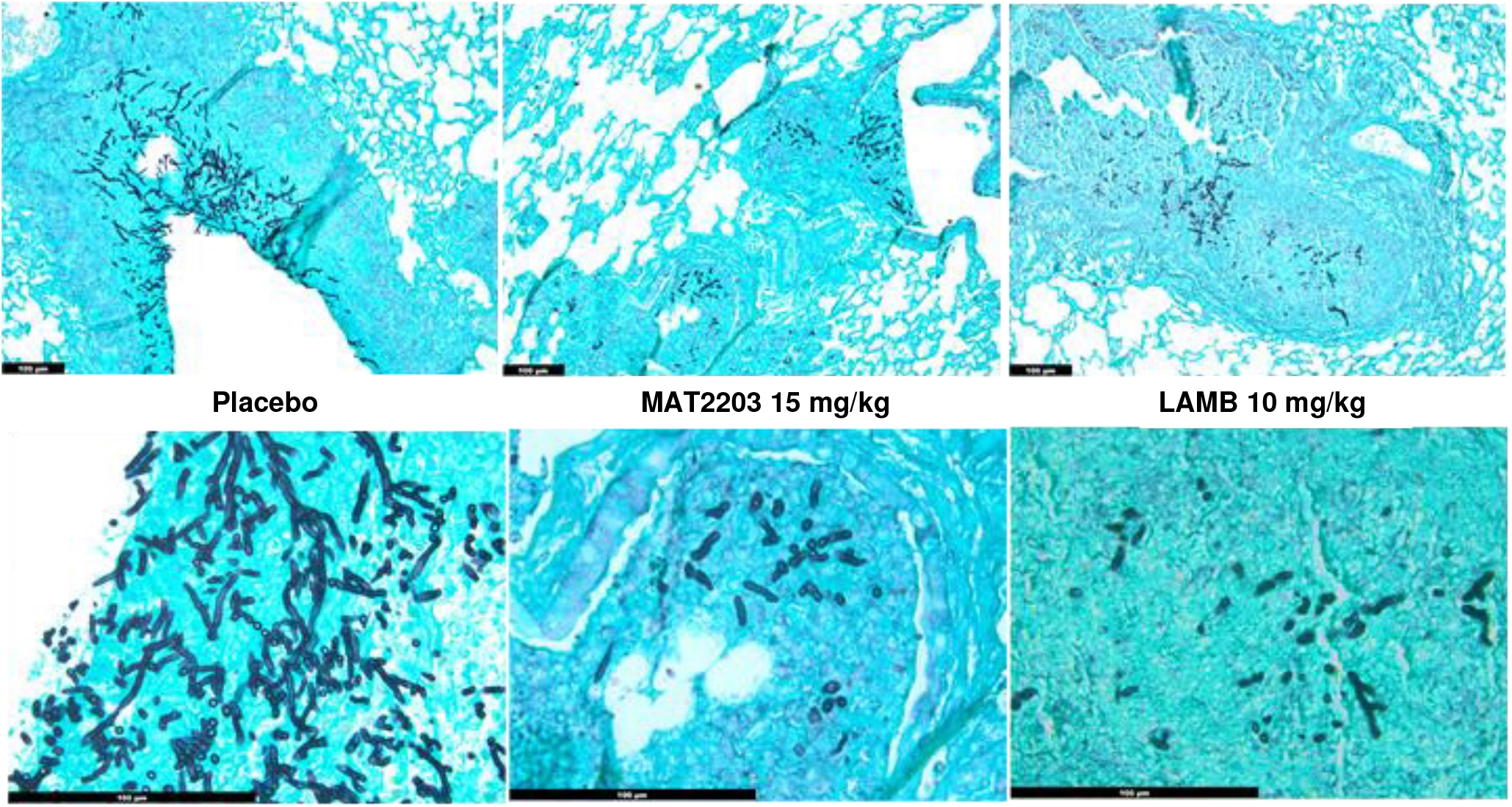
Histology micrographs showing improvement of murine lung infection with MAT2203 or LAMB treatment. Histological examination of lung sections taken from mice infected with *M. circinelloides f. jenssenii* and stained with GMS revealed fungal pneumonia (indicated the abscesses in the placebo mice with broad aseptate hyphae. While evidence of pneumonia still existed in mice treated with either MAT2203 or LAMB, the number of fungal abscesses were less with shorter and damaged fungal hyphae. Bar is 100 µm

## Discussion

In this study, we have shown that MAT2203 has an *in vitro* killing activity that is 5-10-fold higher than LAMB against two clinical isolates of *R. arrhizus* var. *delemar* and *M. circinelloides f. jenssenii* activity (Table 1). The drug demonstrated *in vivo* efficacy in treating *R. delemar or M. circinelloides f. jenssenii* pulmonary infection in immunosuppressed mice. This efficacy is demonstrated by: 1) prolonged median survival time; 2) enhanced overall survival; 3) reduced tissue fungal burden of target organs; and 4) improved histological architecture of infected lungs. Of importance, the *in vivo* efficacy of MAT2203 was equivalent to the efficacy shown by the current standard of care of LAMB.

Initial human clinical studies are consistent with these preclinical study outcomes. In a randomized trial of 141 HIV-positive individuals afflicted by life-threatening cryptococcal meningitis, the oral amphotericin MAT2203 product combined with oral flucytosine resulted in antifungal activity, with similar survival, and less toxicity than intravenous amphotericin B. With six weeks of LNC-enabled oral amphotericin B, statistically fewer lab abnormalities occurred than with one week of intravenous amphotericin B (13). Additionally, in the Matinas’ Compassionate/Expanded Use Access Program, MAT2203 was treated a critically ill 15-year-old female patient, with acute myeloid leukemia and diabetes who suffered from invasive fungal infections in sinus, lung, and brain due to multiple, extremely resistant mucor and aspergillus species. Following only three weeks of therapy on MAT2203, the patient began to show clinical improvement. Renal function returned to normal and repeated sinus/brain MRI showed no evidence of active infection. Additionally, repeated chest CT showed a reduction in pulmonary nodules with no new lesions. The patient continued MAT2203 therapy for a total of 17 weeks with no evidence of nephrotoxicity.

These pre-clinical and clinical efficacy studies, coupled with low toxicity and the additional fact that MAT2203 is administered orally, support continued investigation and development of MAT2203 as a novel, and oral formulation of amphotericin for the treatment of mucormycosis.

## MATERIALS AND METHODS

### Mucorales and culture conditions

*R. arrhizus* var. *delemar* 99-880 and *M. circinelloides f. jenssenii* DI15-131 are clinical isolates obtained from the Fungus Testing Laboratory at the University of Texas Health Sciences Center at San Antonio (UTHSCSA). These two isolates have been used in our murine mucormycosis models (14, 15). The organisms were grown on potato dextrose agar (PDA) plates for 4-7 days until confluent at 37°C. Spores were collected by flooding the plates with sterile phosphate-buffered saline (PBS) containing 0.01% (vol/vol) Tween 80. The spores were concentrated by centrifugation washed in the PBS, diluted, and counted using a hemocytometer. Targeted inoculum for infection was 2.5 × 10^5^ spores/25 μL for *R. arrhizus* var. *delemar* and 2.5 × 10^6^ spores/25 μL for *M. circinelloides f. jenssenii*.

### Susceptibility testing

Mucorales fungi were compared to the activity of LAMB using the (CLSI) M38-A2 method (19).

### Immunosuppression

Male CD-1 mice (20-25 g from Envigo, Indianapolis, IN, USA) were used in this study. Immunosuppression was rendered by administration of cyclophosphamide (200 mg/kg, intraperitoneal [i.p.]) and cortisone acetate (500 mg/kg, subcutaneous [s.c.]) on day -2, +3, and +8 relative to infection. This treatment regimen results ∼ 14 days of leukopenia with total white blood cell count dropping from ∼130000/cm^3^ to almost no detectable leukocytes as determined by Unopette System (Becton-Dickinson and Co.) (14). To prevent bacterial infection, 50 mg/L Baytril (enrofloxacin, Bayer, Leverkusen, Germany) was added to drinking water on Day -3, then switched to daily ceftazidime (5 mg/mouse, s.c.) treatment starting day 0 through day + 13 (20).

### Infection and treatment

Immunosuppressed mice were intratracheally infected with 2.5 × 10^5^ spores of *R. arrhizus* var. *delemar* or 2.5 × 10^6^ spores of *M. circinelloides f. jenssenii* in 25 µL using a gel-loading tip after sedation with isoflurane gas (14). Following inoculation, three mice were sacrificed, and their lungs harvested for quantifying the delivered fungal inoculum by quantitative culturing on PDA plates. Treatment with placebo (diluent control), oral MAT2203 (5 to 45 mg/kg, given once daily [qd] or twice daily [bid] for 7 days) or intravenous injection of 10 mg/kg qd of LAMB (Gilead Sciences Inc., Foster City, CA, USA), started 16 h post infection and continued for 7 days for MAT2203 and 4 days for LAMB. Placebo mice received vehicle control. The primary and secondary endpoints were time to moribundity of infected mice (as determined by the criteria of ruffled and/or matted fur; weight loss of >20%; hypothermia; decreased activity; hunched posture; inability to eat or drink; and torticollis or barrel rolling) and tissue fungal burden in lungs and brains (primary and secondary target organs, respectively) using conidial equivalent by qPCR (21). Histopathological samples were sectioned at 5 micron, and then stained with Grocott methenamine silver (GMS) stain.

### Ethics

Animal studies were approved by the institutional animal care use committee (IACUC) of the Lundquist Institute at Harbor-UCLA Medical Center, according to the NIH guidelines for animal housing and care (approval reference number 22802).

## Statistical analysis

The non-parametric log-rank test was used to determine differences in survival. Differences in tissue fungal burdens were compared by the non-parametric Wilcoxon rank sum test for multiple comparisons. *P* values of < 0.05 were considered significant.

## Acknowledgments

This work was presented at ID Week 2022, Washington, DC (October 21-25) (Submission ID 891240). Research described in this manuscript was conducted at the research facilities of the Lundquist Institute at Harbor-UCLA Medical Center.

## Transparency declarations

A.S.I. served on the Scientific Advisory board of Matinas. T.M., J.C., R.M. are employees of Matinas BioPharma. All other authors have nothing to declare.

